# Rab5 regulates macropinosome closure through recruitment of the inositol 5-phosphatases OCRL/Inpp5b and the hydrolysis of PtdIns(4,5)P_2_

**DOI:** 10.1101/2020.06.08.139436

**Authors:** Michelle E. Maxson, Helen Sarantis, Allen Volchuk, John H. Brumell, Sergio Grinstein

## Abstract

Rab5 is required for macropinosome formation, but its site and mode of action remain unknown. We report that Rab5 acts at the plasma membrane, downstream of ruffling, to promote macropinosome sealing and scission. Dominant-negative Rab5, which obliterates macropinocytosis, had no effect on the development of membrane ruffles. However, Rab5-containing vesicles were recruited to circular membrane ruffles, and SNARE-dependent endomembrane fusion was necessary for completion of macropinocytosis. This fusion event coincided with the disappearance of PtdIns(4,5)P_2_ that accompanies macropinosome closure. Counteracting the depletion of PtdIns(4,5)P_2_ by expression of phosphatidylinositol-4-phosphate 5-kinase impaired macropinosome formation. Importantly, we found that removal of PtdIns(4,5)P_2_ is dependent on Rab5, through the Rab5-mediated recruitment of the inositol 5-phosphatases OCRL and Inpp5b, via APPL1. Knockdown of OCRL and Inpp5b, or APPL1 prevented macropinosome closure, without affecting ruffling. We therefore propose that Rab5 is essential for the clearance of PtdIns(4,5)P_2_ needed to complete macropinosome scission from the plasmalemma.

## Introduction

Macropinocytosis is an actin-driven process that involves the formation and extension of plasma membrane ruffles and the eventual closure of large (≥0.2 to 5.0 µm) endocytic vacuoles (Swanson, 2008). Dendritic cells and macrophages perform macropinocytosis constitutively in order to sample their environment for immune surveillance (Bohdanowicz *et al*, 2013). However, most cell types require stimulation by growth promoters to initiate macropinocytosis (Haigler *et al*, 1979; Mellström *et al*, 1988; Racoosin & Swanson, 1989). In these cases macropinocytosis can play an important role in the delivery of extracellular nutrients, including proteins and amino acids, to lysosomes (Bloomfield & Kay, 2016; Racoosin & Swanson, 1992). The significance of macropinocytosis to cellular physiology is illustrated by its common dysregulation in a variety of diseased states; the manipulation of macropinocytosis is a hallmark of cancer metabolism (O’Donnell *et al*, 2018) and provides an entry gateway for a variety of viruses, including vaccinia and Ebola (Mercer & Helenius, 2012).

The formation of macropinosomes depends on the integration of receptor signalling, phosphoinositide metabolism, activation of small GTPases, and the remodelling of the actin cytoskeleton (Buckley & King, 2017; Levin *et al*, 2014; Marques *et al*, 2017). Growth factors, such as M-CSF and EGF, bind to cognate tyrosine kinase receptors and activate signalling cascades that culminate in the recruitment and activation of phosphatidylinositol 3-kinase (PI3K) (Abella *et al*, 2010; Araki *et al*, 1996; Cheng *et al*, 2015; Wang *et al*, 2001). The resulting generation of phosphatidylinositol-3,4,5-*tris*phosphate (PtdIns(3,4,5)P_3_) is critical for macropinocytosis in a variety of cells types (King & Kay, 2019; Veltman *et al*, 2016; Welliver & Swanson, 2012). Together with its precursor, phosphatidylinositol-4,5-*bis*phosphate (PtdIns(4,5)P_2_), PtdIns(3,4,5)P_3_ promotes the recruitment and activation of small GTPases of the Ras and Rho families that induce the formation of the circular ruffles that precede macropinocytosis (Bar-Sagi & Feramisco, 1986; Fujii *et al*, 2013; Kay *et al*, 2018; Porat-Shliom *et al*, 2008; Welliver *et al*, 2011). Rho-family GTPases support the recruitment of Arp2/3 and formins that mediate the actin polymerization observed in macropinocytic cups (Junemann *et al*, 2016; Veltman *et al*, 2016).

Completion of macropinosome formation requires membrane fusion and is followed by maturation of the resulting vacuole, which proceeds to merge with endosomes and, ultimately, lysosomes. Fusion and maturation are accompanied by rapid disassembly of the rich actin meshwork that initiated ruffling and macropinosome formation. The disassembly is due, at least in part, to the breakdown of PtdIns(3,4,5)P_3_ and PtdIns(4,5)P_2_ that accompanies macropinosome closure (Araki *et al*, 2007; Hasegawa *et al*, 2011; Maekawa *et al*, 2014; Welliver & Swanson, 2012; Yoshida *et al*, 2009). Closure is also coincident with the acquisition of small GTPases of the Rab family, notably Rab5 (Araki *et al*, 2006; Feliciano *et al*, 2011; Porat-Shliom *et al*, 2008; Roberts *et al*, 2000; Welliver & Swanson, 2012). Rab5, a prototypical component of early endosomes, is thought to drive the initial stages of macropinosome maturation. However, expression of dominant-negative Rab5 was found to block macropinocytosis (Feliciano *et al*, 2011; Roberts *et al*, 2000), suggesting an additional, early role for Rab5 in macropinosome formation. Accordingly, active Rab5 has been localized to the plasma membrane during macropinocytosis (Bucci *et al*, 1992; Chavrier *et al*, 1990; Lanzetti *et al*, 2004). At this time, however, the manner whereby Rab5 affects macropinosome formation remains unclear. We therefore revisited the role of Rab5 in macropinocytosis.

## Results

### Rab5 is necessary for the completion of macropinocytosis

Serum-starved A431 cells respond quickly to the addition of EGF with plasmalemmal ruffling and the formation of 1.5-2 μm-sized macropinosomes (Fabricant *et al*, 1977; Haigler *et al*, 1979). We chose this well-established model system to investigate the role of Rab5 in macropinocytosis. As expected, in the absence of EGF A431 cells internalized little tetramethylrhodamine-conjugated 70 kDa dextran (TMR-D), a prototypical marker of macropinocytic uptake, and uptake was unchanged by expression of wild type (WT) Rab5A (Fig 1A, left panel). After incubation with EGF, Rab5A-positive cells internalized TMR-D robustly (Fig 1A, right panel), as did non-transfected neighbouring cells (not shown). In contrast, EGF-treated A431 cells expressing a dominant-negative (DN) version of Rab5A (Rab5A_S34N_) failed to internalize TMR-D, while the non-transfected neighbouring cells did (Fig 1B). Expression of Rab5A_S34N_ decreased macropinocytosis by 86.8% (Fig 1C, black bars), in agreement with earlier findings (Feliciano *et al*, 2011; Roberts *et al*, 2000). Because expression of dominant-negative alleles can have pleiotropic effects, e.g. by inhibiting exchange factors shared by multiple GTPases, we validated the requirement for Rab5 by silencing the three Rab5 isoforms; the combined use of siRNA to the A, B and C isoforms decreased the overall expression of Rab5 by 88.1 ± 4.3% (n= 4) as assessed by immunoblotting (not shown). This collective silencing depressed TMR-D uptake by 73.0% (Fig 1C, grey bars), validating the specific involvement of Rab5 in macropinosome formation.

**Figure 1.**
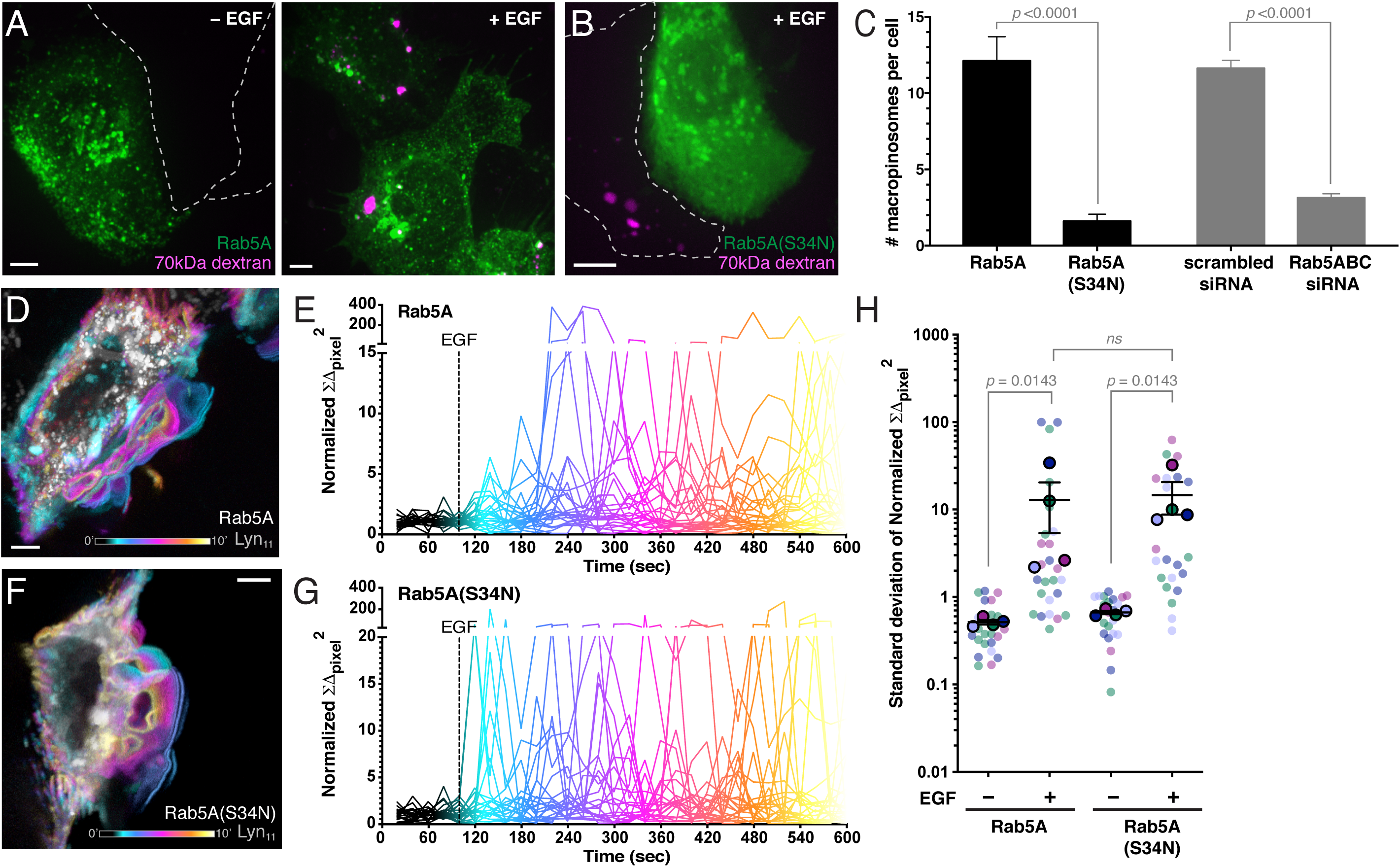
Rab5 is required for the completion of macropinocytosis, but not for cell ruffling. Macropinocytosis was induced in A431 cells by EGF (see Materials and Methods). **A)** Effect of Rab5A overexpression. Extended focus visualization of TMR-D uptake by GFP-Rab5A-expressing cells in the absence (left) or presence (right) of EGF. TMR-D is shown in magenta. In this and subsequent figures, the outlines of untransfected A431 cells are indicated by dotted lines. **B)** Effect of Rab5A_S34N_ overexpression. Extended focus visualization of TMR-D uptake by GFP-Rab5A_S34N_-expressing cells stimulated by EGF. **C)** After EGF treatment in the presence of TMR-D, transiently transfected or siRNA-treated A431 monolayers visualized and the number of macropinosomes per cell counted by confocal microscopy. For each condition, ≥5 independent experiments were quantified, with ≥ 10 cells per replicate. *p* calculated using unpaired, 2-tailed students t-tests. Data are means ± SEM. **D)** Temporal projection of cellular ruffling Rab5A-expressing cells over a 10 min incubation with EGF. GFP-Rab5A is pseudo-colored white, Lyn_11_-RFP is pseudo-colored as a function of time using the “cool” LUT. Color-coded temporal scale is included. **E)** Quantitation of ruffling for Rab5A-expressing cells over a 10 min timespan. Individual cells are represented by separate traces, *N* = 23. Lines are temporally pseudo-colored as in **D**. Addition of EGF is indicated by dotted vertical line. Detailed description of normalized ΣΔ_pixel_^2^ calculation can be found in Materials and Methods. **F)** Temporal projection of cellular ruffling Rab5A_S34N_-expressing cell stimulated with EGF as in **D**. Images in **A-F** are representative of ≥ 30 fields from ≥ 3 separate experiments of each type. All scale bars: 5 μm. **G)** Quantitation of ruffling for Rab5A_S34N_-expressing cells, as in **E**; *N* = 22. **H)** Scatter plot of standard deviations of the normalized ΣΔ_pixel_^2^ values in the experiments illustrated in **E** and **F**. Rimmed circles indicate means of individual experiments, identified by color-coding. Means for 4 replicates ± SEM are shown. *p* values calculated using the Mann-Whitney test.

Macropinocytosis entails the development of membrane ruffles and their closure to form sealed vacuoles. We therefore sought to define whether Rab5 is required for the extension of ruffles or, instead, for their sealing. To this end, the dynamics of macropinosome formation was monitored by time-lapse video microscopy, utilizing the N-terminal domain of Lyn (Lyn_11_) tagged with RFP to selectively visualize the plasma membrane. This region of Lyn is myristoylated and palmitoylated, effectively targeting the Lyn_11_-RFP construct to the plasma membrane and nascent macropinosomes. Temporal projections of A431 cells expressing Lyn_11_-RFP revealed the extensive ruffling elicited by treatment with EGF (e.g. Fig 1D). Importantly ruffling was qualitatively indistinguishable in cells that were co-transfected with Lyn_11_-RFP and either WT Rab5A or Rab5A_S34N_ (cf. Fig 1D and 1F; cf. also Appendix Movie 1 and 2). A more rigorous comparison of the effects of the WT and DN forms of Rab5 was made by measuring the sum of the differences in fluorescence intensity (ΣΔ_pixel_) associated with membrane displacement (i.e. ruffling). The differences were squared (ΣΔ_pixel_^2^) to eliminate the cancelling effects of positive vs. negative changes in fluorescence intensity. As shown in Fig 1E and G, where the behaviour of multiple individual cells is compared, membrane positioning changed comparatively little over time prior to addition of EGF, but marked oscillations were apparent upon addition of the growth factor. Comparison of the standard deviation of the ΣΔ_pixel_^2^ in the period before and after addition of EGF confirmed that the ruffling triggered by the growth factor was similar in cells expressing Rab5A or Rab5A_S34N_ (Fig 1H). We therefore concluded that functional Rab5 is not required for membrane ruffling and that the GTPase is instead involved in another, downstream step in vacuole generation.

### Rab5 is recruited to plasmalemmal circular ruffles prior to macropinosome closure

Because Rab5 is seemingly not required for ruffling, we considered the possibility that it is instead involved in macropinosome sealing. This would require engagement of the GTPase prior to closure of the macropinocytic vacuole. Rab5 has been reported to be present in macropinosome membranes shortly after closure (Feliciano *et al*, 2011; Roberts *et al*, 2000), but the precise timing of its acquisition has not been established. In accordance with the earlier reports, we routinely observed Rab5 merging with nascent macropinosomes within ≈4 min of EGF addition (Fig 2A). While examining the macropinocytic process by time-lapse microscopy we observed the striking recruitment and tight apposition of GFP-Rab5A-positive endocytic vesicles to emerging circular ruffles/early macropinosomes as early as ≈1 min after EGF addition (Fig 2A and B; Appendix Movie 3).

**Figure 2.**
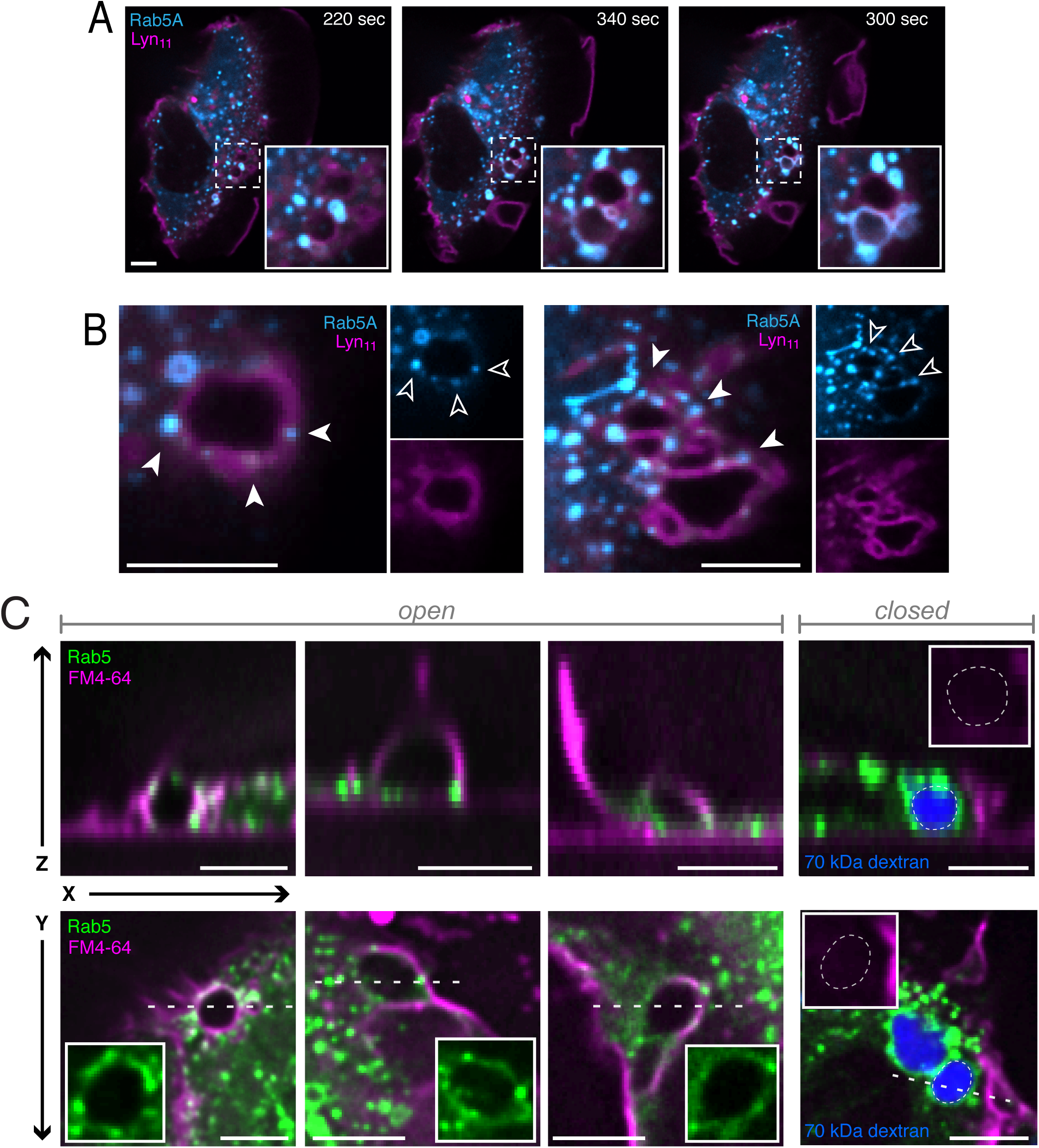
Rab5A is recruited to circular ruffles and open macropinocytic cups. **A)** Rab5A localization to nascent macropinosomes. Confocal visualization of cells expressing GFP-Rab5A (cyan) and Lyn_11_-RFP (magenta), treated with EGF to induce macropinocytosis. The time after stimulation when images were acquired is indicated. Insets are 2.8x magnifications of area in dotted square. **B)** Localization of GFP-Rab5A (cyan) in plasma membrane ruffles visualized with Lyn_11_-RFP (magenta) during EGF-induced macropinocytosis. Arrowheads mark Rab5-positive vesicles apposed to ruffles. Insets show individual Rab5A and Lyn_11_ channels. **C)** Macropinocytosis was induced by EGF in A431 cells expressing GFP-Rab5A, and after 2 min cells were cooled and visualized by confocal microscopy. Surface membranes and open macropinosomes were visualized by FM4-64 staining (see Materials and Methods). The localization of Rab5A (green) and FM4-64 in *XZ* reconstructions (top) and individual *XY* optical slices (bottom) are shown for 3 representative open macropinosomes and 1 closed macropinosome, as indicated. Closed macropinosomes retained TMR-D, shown in blue. Dotted line across *XY* slice represents location of corresponding *XZ* reconstruction. Insets are 1.6, 1.1, 1.2 and 1.3x magnification (left to right).

When acquiring optical slices by confocal microscopy it is not possible to conclusively differentiate between open circular ruffles and sealed vacuoles. This strategy alone was therefore insufficient to establish whether the tight apposition and fusion of Rab5 vesicles occur prior to macropinosome sealing. To more definitively establish the timing of Rab5 recruitment, we induced macropinocytosis with EGF for 2 min in the presence of TMR-D, then rapidly chilled the cells to arrest the process and stained them with the membrane-impermeant dye FM4-64. FM4-64, which is brightly fluorescent only when intercalated into membranes, was viewed immediately (< 5 min) to prevent its endocytic uptake. Thus, only the plasma membrane –which is continuously exposed to the extracellular milieu– as well as open (but not sealed) macropinosomes, were expected to show dye fluorescence. Conversely, only sealed macropinosomes were expected to retain TMR-D. Strikingly, we found that Rab5A-positive vesicles were distinctly recruited to circular ruffles that had not closed, identified as round structures that were nearly continuous yet stained clearly with FM4-64; multiple examples are illustrated in transverse (*X* vs *Y*; bottom) and in orthogonal reconstructions (*X* vs *Z*; top) in Fig 2C. These structures did not retain TMR-D (Fig 2C, left), which was instead retained by FM4-64-negative macropinosomes (Fig 2C, right). Although we cannot ascertain that the recruited endosomes had fused with open circular ruffles, these data indicate that tight apposition of Rab5-positive organelles precedes macropinosome closure.

### Rab5 plays a role at the plasma membrane promoting the completion of macropinocytosis

Because Rab5-containing vesicles were recruited to circular ruffles at early stages of macropinosome formation (Fig 2), and genetic inhibition of Rab5 blocked macropinocytosis (Fig 1B-C), we questioned whether their attachment and/or fusion with the plasma membrane is\ required to promote macropinosome formation. To this end, we initially utilized an inducible rapamycin heterodimerization system (iRAP) to recruit WT Rab5A or Rab5A_S34N_ to the plasma membrane. Cells were transfected with CFP-FKBP-Rab5A or CFP-FKBP-Rab5A_S34N_, plus a plasma membrane-localizing Lyn_11_-FKB construct; incubation of the cells with rapamycin resulted in the rapid (≈1 min) recruitment of Rab5A or Rab5A_S34N_ to the plasma membrane (Fig 3A and C insets). Recruitment of WT Rab5A to the plasma membrane was neither sufficient to induce macropinocytosis –measured as TMR-D internalization– nor did it prevent that induced by EGF (Fig 3A and B). In contrast, near complete recruitment of Rab5A_S34N_ to the plasma membrane was still able to robustly inhibit macropinocytosis (Fig 3C and D), in a manner indistinguishable from that of Rab5A_S34N_ expressed in the cytosol, which is in equilibrium with membrane-associated Rab5A_S34N_ by virtue of its geranylgeranylation (Fig 1B and 3D). These observations suggest that the dominant negative effect of Rab5A_S34N_ on macropinocytosis is exerted locally at the plasma membrane, a consequence of inhibited Rab5 function during presumptive plasmalemmal association.

**Figure 3.**
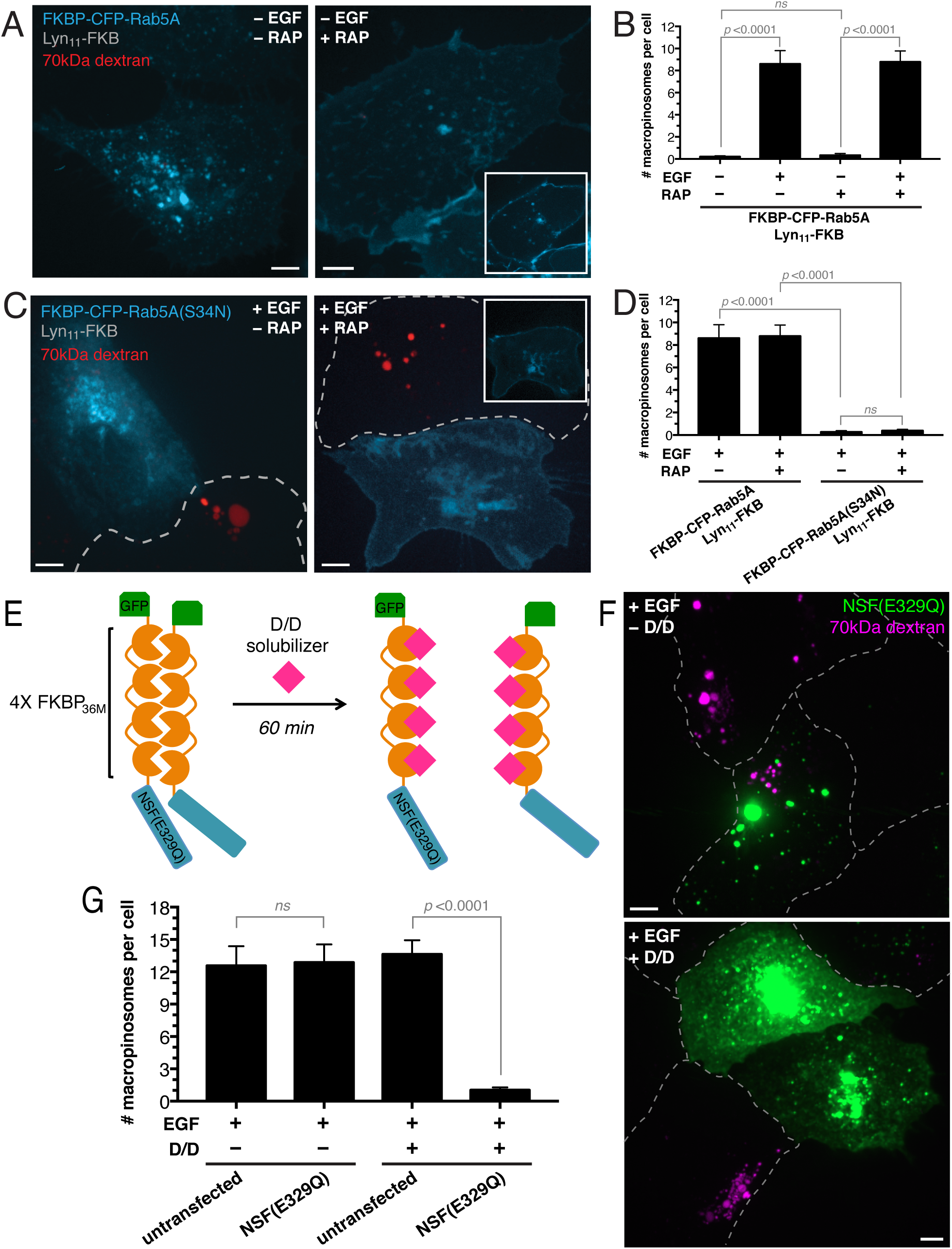
Rab5A localization to the plasma membrane, through SNARE mediated fusion events, is necessary for completion of macropinocytosis. **A)** Cells were transfected with FKBP-CFP-Rab5A (cyan) and Lyn_11_-FKB and incubated with TMR-D (red) in the absence of EGF. In the experiment shown on the right, recruitment of Rab5A to the plasma membrane was induced by rapamycin. Main panels are extended focus images. Inset: single *XY* optical slice illustrating plasma membrane-recruited Rab5A. **B)** Effect of Rab5A recruitment on macropinocytosis. Cells transfected with FKBP-CFP-Rab5A and Lyn_11_-FKB were treated with or without rapamycin, in the presence or absence of EGF plus TMR-D, as indicated. The number of macropinosomes per cell was counted as described in Materials and Methods. For each condition, 3 independent experiments were quantified, with ≥ 10 cells per replicate. Data are means ± SEM. *p* values calculated using unpaired, 2-tailed student’s t-tests. **C)** Cells were transfected with FKBP-CFP-Rab5A_S34N_ (cyan) and Lyn_11_-FKB and incubated with TMR-D (red) in the presence of EGF. Where indicated (right) Rab5A _S34N_ recruitment to the plasma membrane was induced by rapamycin. Other details as in **A. D)** Effect of Rab5A or Rab5A_S34N_ recruitment on macropinocytosis. Cells transfected with FKBP-CFP-Rab5A/Rab5A_S34N_ and Lyn_11_-FKB were treated with or without rapamycin, in the presence of EGF plus TMR-D, as indicated. Other details as in **B.** For each condition, 3 independent experiments were quantified, with ≥ 10 cells per replicate. *p* value was calculated using the unpaired, 2-tailed students t-test. Data are means ± SEM. Note that data Rab5 plus EGF in **D** are replotted from **B. E)** Schematic representation of the reverse aggregation of the F_M_4-NSF_E329Q_ construct tagged with EGFP. When expressed in mammalian cells, EGFP-F_M_4-NSF_E329Q_ aggregates spontaneously but aggregation is reversed upon addition of D/D solubilizer, allowing for acute inhibition of SNARE-mediated vesicle fusion. **F)** Cells expressing EGFP-F_M_4-NSF_E329Q_ were incubated in the absence (top) or 60 min after addition of D/D solubilizer. Cells were then treated with EGF plus TMR-D, as described in Materials and Methods. Following fixation, transfected cells and untransfected neighbours were monitored for retention of TMR-D (magenta). All scale bars: 5 μm. Images in **A,C** and **F** are representative of ≥ 30 fields from ≥ 3 separate experiments of each type. **G)** Cells treated as in **F** were fixed and the number of macropinosomes per cell counted by confocal microscopy in transfected cells and untransfected neighbors. For each condition, 3 independent experiments were quantified, with ≥ 10 cells per replicate. Data are means ± SEM. *p* value was calculated using the unpaired, 2-tailed students t-test.

Inactive, GDP-bound Rab5 is thought to reside in the cytosol, complexed with Rab GDI, while the active GTP-bound form associates with early endocytic membranes (Zerial & McBride, 2001). During macropinocytosis, it was conceivable that Rab5 could be activated/recruited to the plasma membrane from the cytosol, or be acquired through fusion of endocytic vesicles containing active Rab5, as suggested by the observations in Fig 2. To distinguish between these mechanisms, we adapted a reverse dimerization system (Rivera *et al*, 2000) to acutely block SNARE-mediated membrane fusion with dominant negative N-ethylmaleimide-sensitive factor (NSF_E329Q_). Expression of this reverse dimerization construct (EGFP-F_M_4-NSF_E329Q_) results in formation of multimeric aggregates in the cytosol –due to the presence of four tandem mutant FKBP (F_M_4) domains in the construct– which can be solubilized promptly by the addition of a cell-permeant rapamycin analog (Fig 3E). Due to the acute release of NSF_E329Q_, the cells do not experience the toxicity normally seen when expressing soluble dominant negative NSF for extended periods (Coppolino *et al*, 2001).

As expected, when expressed in A431 cells EGFP-F_M_4-NSF_E329Q_ formed large cytosolic aggregates (Fig 3F, top) that were functionally inconsequential. This was verified by monitoring the appearance of the Golgi complex, which was unaffected by the aggregated form of the dominant negative NSF (Appendix Fig S1A). As expected, aggregated EGFP-F_M_4-NSF_E329Q_ was also without effect on macropinocytosis (Fig 3F-G). The EGFP-F_M_4-NSF_E329Q_ aggregates dissociated gradually upon addition of the D/D solubilizer (Fig 3E-F). The appearance of soluble (EGFP-F_M_4-NSF_E329Q_ was accompanied by inhibition of endomembrane fusion, as evidenced by the fragmentation and dispersal of the Golgi complex (Appendix Fig S1A, right). Importantly, solubilization of the dominant-negative NSF caused a profound (92%) inhibition of macropinocytosis (Fig 3F-G). Of note, the membrane ruffling induced by EGF was unaffected by the soluble NSF_E329Q_ (Appendix Fig S1B). We therefore concluded that a SNARE-mediated fusion event, likely that of Rab5-containing vesicles with the plasma membrane, is required for macropinocytosis.

### Rab5 contributes to the depletion of PtdIns(4,5)P_2_ at the macropinocytic cup, which is required for macropinosome sealing

Loss of PtdIns(4,5)P_2_, which is abundant in the plasma membrane, is one of the early events that accompany macropinosome formation (Porat-Shliom *et al*, 2008; Welliver & Swanson, 2012). While examining EGF-induced macropinocytosis by time-lapse microscopy we observed that the docking of Rab5A-positive endocytic vesicles onto circular ruffles preceded the loss of PtdIns(4,5)P_2_ (Fig 4A). Strikingly, expression of dominant negative Rab5A_S34N_ prolonged the presence of PtdIns(4,5)P_2_ on circular ruffles (Fig 4B), suggesting that active Rab5 is normally required for the depletion of the inositide. These observations suggested that PtdIns(4,5)P_2_ depletion may be required for completion of macropinocytosis.

**Figure 4.**
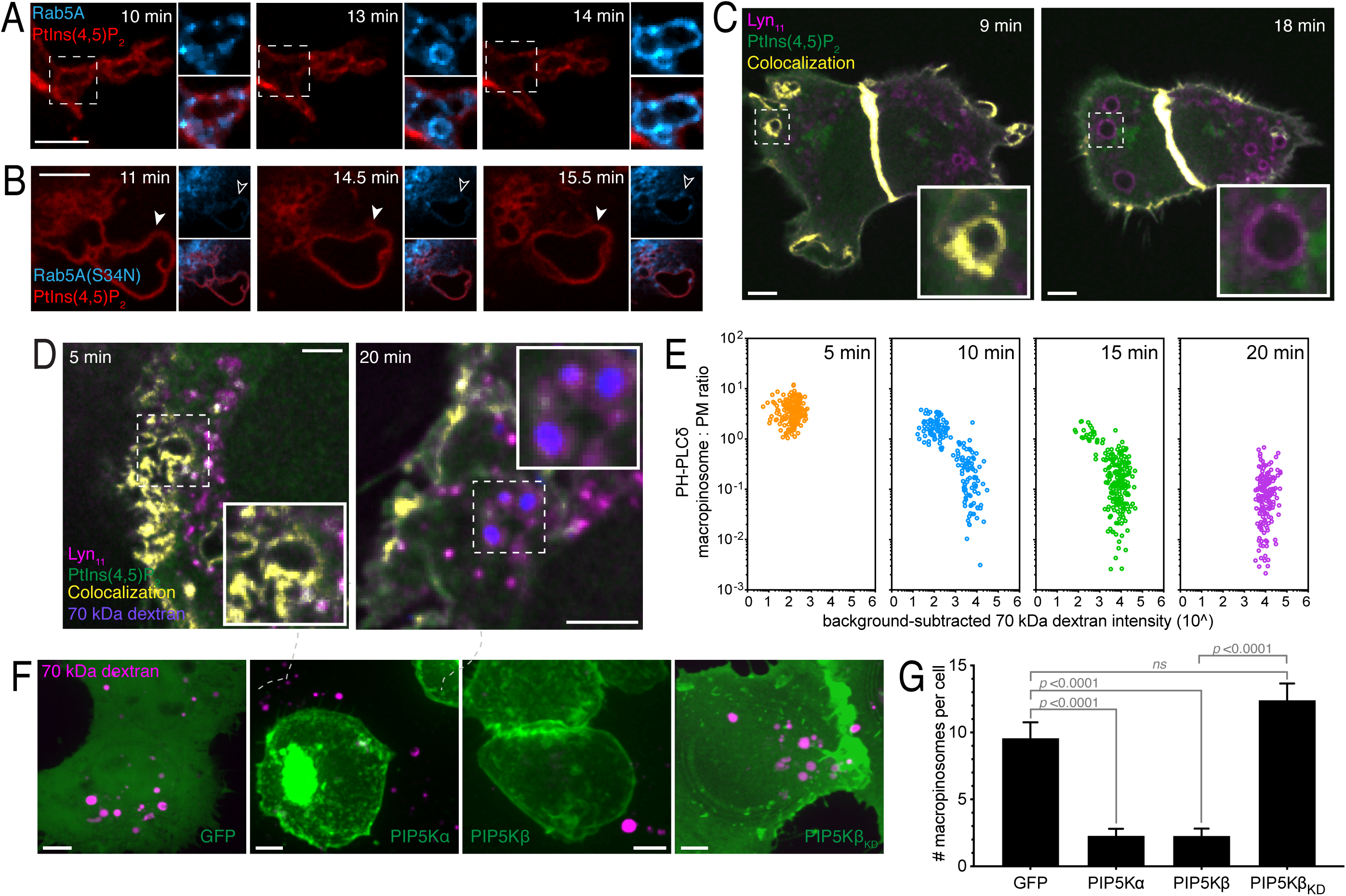
PtdIns(4,5)P_2_ disappearance from forming macropinosomes is necessary for macropinosome closure. **A)** Live cell visualization of PtdIns(4,5)P_2_ using PLCδ-PH-RFP (red) in cells co-expressing GFP-Rab5A (cyan), at indicated times after EGF addition. Panel insets: Rab5A and merged channels, at 2x magnification. **B)** Visualization of PLCδ-PH-RFP (red) in cells co-expressing GFP-Rab5A_S34N_ (cyan), at indicated times after EGF addition. Panel insets: Rab5A_S34N_ and merged channels. **C)** Live cell visualization of PtdIns(4,5)P_2_ using PLCδ-PH-GFP (green) and the plasma membrane using Lyn_11_-RFP (magenta) in EGF-treated cells, at indicated timepoints. EGF addition occurred between the 150-180 sec timepoints. Colocalization of PLCδ-PH-GFP with Lyn_11_-RFP shown in yellow. Insets: 4x magnification. **D)** Cells transfected with PLCδ-PH-GFP (green) and Lyn11-RFP (magenta) were treated with EGF in the presence of TMR-D, as described in Materials and Methods. Retention of TMR-D (blue) was visualized by fixing cells 5 min or 20 min after EGF addition followed by serial confocal imaging and extended focus projection. Colocalization of PLCδ-PH-GFP with Lyn_11_-RFP shown in yellow. Insets: 1.7x magnification. **E)** Plots of PLCδ-PH-GFP macropinosome:plasma membrane ratio vs background-subtracted TMR-D intensity for all Lyn_11_-RFP-positive macropinocytic circular ruffles/vacuoles, determined at 5, 10, 15 and 20 min after EGF addition. Number of vacuoles quantified: 5 min, 177; 10 min, 184; 15 min, 225; 20 min, 179. **F)** Monolayers transfected with GFP, GFP-PIP5Kα, GFP-PIP5Kβ or GFP-PIP5Kβ_KD_ (all in green) were treated with EGF in the presence of TMR-D, as described in Materials and Methods. Following fixation, retention of TMR-D (magenta) was imaged as above. Images in **A-F** are representative of ≥ 30 fields from ≥ 3 separate experiments of each type. All scale bars: 5 μm. **G)** Cells treated as in **F** were fixed and the number of macropinosomes per cell counted by confocal microscopy. For each condition, 3 independent experiments were quantified, with ≥ 10 cells per replicate. Data are means ± SEM. *p* values calculated using unpaired, 2-tailed student’s t-tests.

This notion was tested by expressing the genetically-encoded PtdIns(4,5)P_2_ biosensor PLCδ-PH-GFP in cells also expressing Lyn_11_-RFP, used to visualize the membrane and nascent macropinosomes. As expected, PtdIns(4,5)P_2_ was present in the plasma membrane and was clearly visible in the ruffles induced by EGF (Fig 4C and Appendix Movie 4). As reported (Porat-Shliom *et al*, 2008; Welliver & Swanson, 2012), PtdIns(4,5)P_2_ was lost from the macropinosome membrane as it internalized (Fig 4C, right and Appendix Movie 4). The loss of PtdIns(4,5)P_2_ from the macropinosome membrane was associated with the retention of TMR-D, the hallmark of closed macropinosomes (Fig 4D). This inverse correlation was analyzed in detail measuring the abundance of PLCδ-PH-GFP on Lyn_11_-RFP-positive cups/vesicles over time and comparing it to the retention of TMR-D. Numerous circular ruffles containing high PtdIns(4,5)P_2_ and little TMR-D were observed 5 min after stimulation with EGF (Fig 4E). Between 10 and 20 min the number of vacuoles retaining TMR-D increased progressively, as the PtdIns(4,5)P_2_ levels decreased (Fig 4E; Appendix Fig S2A). Analysis of these populations (Appendix Fig S2B) showed the vesicles fell into well-defined groups: those that were open (i.e. devoid of TMR-D) and contained PtdIns(4,5)P_2_, and those that were seemingly closed (i.e. able to retain TMR-D) and depleted of PtdIns(4,5)P_2_. There were virtually no sealed vesicles that retained PtdIns(4,5)P_2_. Taken together, these data confirmed that PtdIns(4,5)P_2_ localized predominantly to open macropinocytic cups, and was depleted quickly at the time of macropinosome sealing.

These observations raised the possibility that PtdIns(4,5)P_2_ depletion is in fact required for the completion of macropinocytosis. To examine this possibility, we sought to prevent this depletion by overexpressing the type I PtdIns 4-phosphate 5-kinase (PIP5K), which catalyzes the conversion of plasmalemmal PtdIns(4)P to PtdIns(4,5)P_2_. When expressed heterologously in A431 cells, the GFP-tagged α and β isoforms of PIP5K localized to the plasma membrane, where they effectively blocked macropinocytic retention of TMR-D by 76.2% and 76.3%, respectively, compared to cells expressing GFP only (Fig 4F-G). Notably, expression of a kinase-dead version of PIP5Kβ that is targeted normally to the membrane did not block macropinocytosis, indicating that the catalytic activity of PIP5K, and thus the overproduction of PtdIns(4,5)P_2_, is responsible for blocking the completion of macropinocytosis. We interpreted this to mean that sustained presence of PtdIns(4,5)P_2_ blocks macropinosome sealing.

### Rab5 effectors Inpp5b, OCRL and APPL1 localize to macropinocytic cups and vesicles and are required for macropinosome sealing

The timing of Rab5 acquisition and loss of PtdIns(4,5)P_2_ from forming phagosomes led us to question whether the two events are causally related. The mammalian 5-phosphatases Inpp5b and OCRL, which can degrade PtdIns(4,5)P_2_, are both Rab5-associating effectors implicated in endocytosis and macropinocytosis (Fukuda *et al*, 2008; Hyvola *et al*, 2006; Mao *et al*, 2009; Shin *et al*, 2005; Swan *et al*, 2010; Williams *et al*, 2007). These phosphatases were therefore likely candidates to catalyze the observed PtdIns(4,5)P_2_ depletion. Inpp5b and OCRL can interact with Rab5 directly, or interact with endomembranes by way of another Rab5 effector, APPL1 (Bohdanowicz *et al*, 2011; Erdmann *et al*, 2007; Swan *et al*, 2010; Zhu *et al*, 2007). We observed that soon after stimulation by EGF, punctate structures containing Inpp5b and OCRL –likely endomembrane vesicles– were recruited to PtdIns(4,5)P_2_-positive plasma membrane ruffles (Fig 5A), in a manner similar to that observed for Rab5. Notably, APPL1 was also recruited in this fashion (Fig 5A, right). Time-lapse imaging revealed that the recruitment of Inpp5b, OCRL and APPL1 preceded the loss of PtdIns(4,5)P_2_ from macropinocytic cups (Appendix Fig S3). Like the analogous early recruitment of Rab5-positive vesicles, we interpreted this as evidence that the docking of Inpp5b-, OCRL- and APPL1-bearing vesicles occurred prior to macropinosomes sealing. Extensive acquisition of Inpp5b, OCRL and APPL1 was obvious on macropinosomes that retained TMR-D, that were therefore closed (Fig 5B), consistent with earlier findings (Swan *et al*, 2010; Wall *et al*, 2018; Zoncu *et al*, 2009).

**Figure 5.**
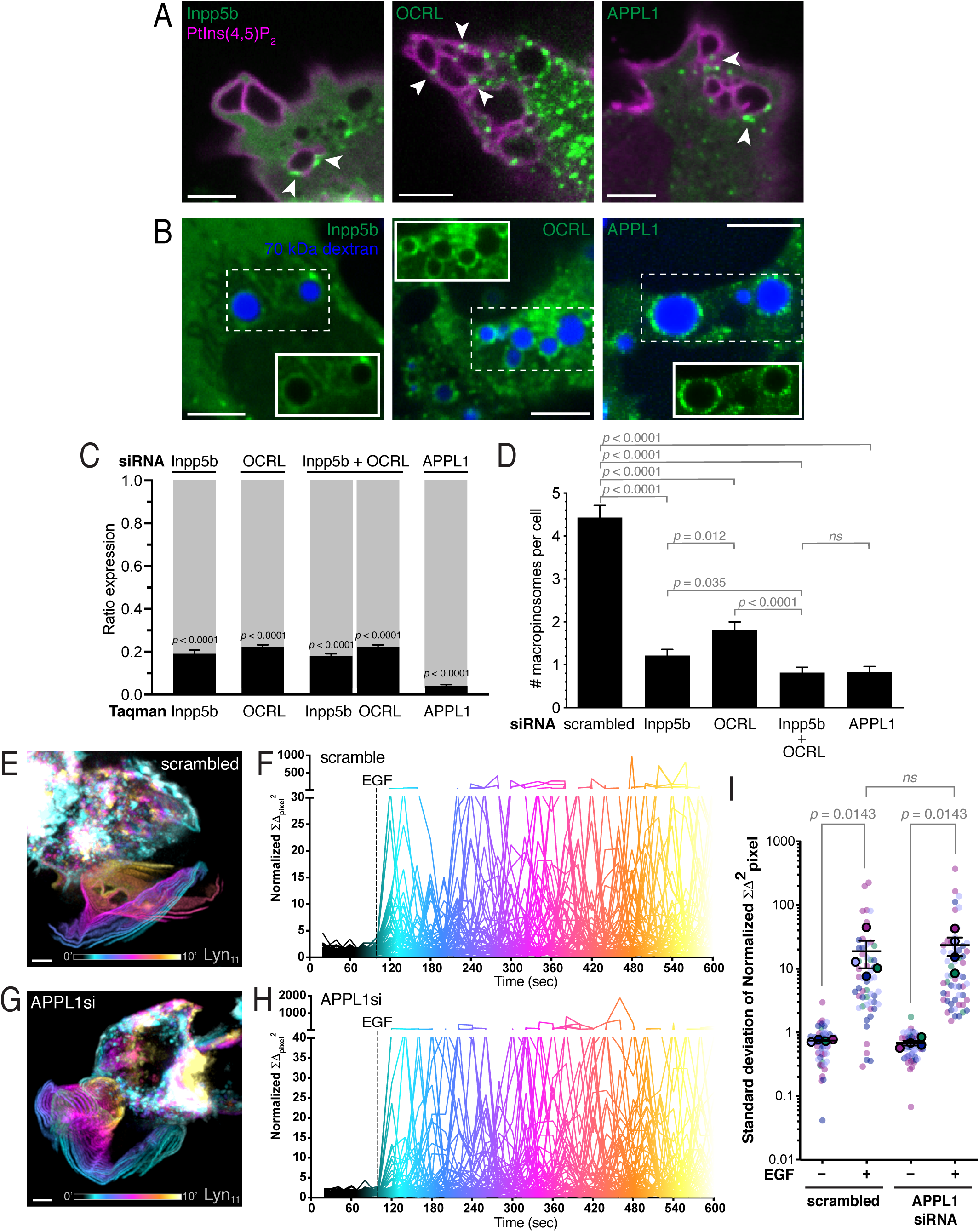
Role of Inpp5b, OCRL and APPL1 in macropinocytosis. **A)** Macropinocytosis was induced by EGF addition in cells expressing GFP-tagged Inpp5b, OCRL or APPL1 (green), with PLCδ-PH-RFP (magenta). Arrowheads indicate apposition of Inpp5b, OCRL or APPL1-positive vesicles onto PtdIns(4,5)P_2_-positive macropinocytic ruffles/vacuoles. **B)** Monolayers transfected with GFP-tagged Inpp5b, OCRL or APPL1 (green) were treated EGF in the presence of TMR-D, as described in Materials and Methods. Following fixation, retention of TMR-D was imaged (blue). Insets: GFP-tagged Inpp5b, OCRL or APPL1 channels alone. **C)** Effect of siRNA-mediated gene silencing, singly or in combination as indicated, on the endogenous expression of Inpp5b, OCRL or APPL1. Expression was measured after 48 hr treatment using qPCR and the ΔΔ*C*_*T*_ method. Data are expressed as fractional expression (black) relative to expression in cells treated with scrambled siRNA (grey). **D)** Effect of siRNA-mediated gene silencing of Inpp5b, OCRL, Inpp5b+OCRL, or APPL1 on macropinocytosis. Monolayers were treated with indicated siRNAs for 48 hr, then treated EGF in the presence of TMR-D. Macropinosome formation was scored as described in Materials and Methods. For each condition, 3 independent experiments were quantified, with ≥ 10 cells per replicate. Data are means ± SEM. *p* values calculated using unpaired, 2-tailed student’s t-tests. **E)** Temporal projection of cellular ruffling in Lyn_11_-RFP-expressing cells treated with control (scrambled) siRNA. Cells were stimulated with EGF and ruffling visualized over 10 min. See Fig 1D for details. **F)** Quantitation of ruffling in cells treated with control (scrambled) siRNA. *N* = 48. Addition of EGF indicated by dotted vertical line. See Fig 1E for details. **G)** Temporal projection of cellular ruffling in Lyn_11_-RFP-expressing cells treated with APPL1 siRNA. Cells were stimulated with EGF and ruffling visualized over 10 min. Scale bars in **A-G**: 5 μm. Images are representative of ≥ 30 fields from ≥ 3 separate experiments of each type. **H)** Quantitation of ruffling in cells treated with APPL1 siRNA. *N* = 53. **I)** Scatter plot of standard deviations of the normalized ΣΔ_pixel_^2^ values in the experiments illustrated in **F** and **H**. Rimmed circles indicate means of individual experiments, identified by color-coding. Means for 4 replicates ± SEM are shown. *p* values calculated using the Mann-Whitney test.

The functional requirement of these proteins for macropinocytosis was assessed by RNA-mediated silencing. The 5-phosphatases were silenced individually or together; APPL1 was silenced as well. The effectiveness of the silencing protocol was assessed first: the expression of Inpp5b was reduced by 80.9% and that of OCRL by 77.8%, when silenced separately. When silenced jointly expression was suppressed by 82.1% and 77.7% for Inpp5b and OCRL, respectively. APPL1 silencing was most effective, reaching 95.9% (Fig 5C). The effects of these treatments on macropinocytosis were assessed next. Inhibiting the expression of either Inpp5b or OCRL caused a marked reduction in macropinocytic efficiency, while simultaneous silencing of both phosphatases resulted in an even greater reduction (≈85%; Fig 5D). The extent of the latter was comparable to the one attained by silencing APPL1 (Fig 5D).

These findings are consistent with the notion that APPL1 cooperates with Rab5 in the recruitment of OCRL and Inpp5b. We therefore utilized APPL1-silenced cells to further assess the role of PtdIns(4,5)P_2_ breakdown by the phosphatases in the macropinocytic process. The effect of down-regulating the adaptor on membrane ruffling was investigated, using the same approaches introduced earlier. Visualization (Fig 5E vs. G; compare also Appendix Movie 5 vs. 6) and quantitative analysis (Fig F, H and I) of ruffling in cells transfected with Lyn_11_-RFP showed no significant differences between cells treated with scrambled siRNA and those silenced using APPL1-directed siRNA, which responded normally to stimulation by EGF. These findings reinforce the conclusion that depletion of PtdIns(4,5)P_2_ occurs after plasma membrane ruffling and is likely involved in macropinosome sealing.

### Recruitment of yeast Inp54 to the plasma membrane bypasses the dominant-negative effect of Rab5A_S34N_ on macropinocytosis

Jointly, the preceding observations suggest that Rab5 may participate in macropinosome sealing by recruiting Inpp5b and OCRL, which facilitate sealing by dephosphorylation of PtdIns(4,5)P_2_. If this model were true, we hypothesized that it would be possible to bypass the inhibitory effect of Rab5A_S34N_ by recruiting to the plasma membrane an alternative, heterologous 5-phosphatase capable of hydrolyzing PtdIns(4,5)P_2_ while the cells are undergoing EGF-induced ruffling. We chose the yeast 5-phosphatase Inp54, since it is exquisitely specific for PtdIns(4,5)P_2_ (Raucher *et al*, 2000; Stolz *et al*, 1998). As before, we utilized an iRAP system to recruit Inp54 to the plasma membrane of cells expressing either Rab5A or Rab5A_S34N_. These cells were co-transfected with YFP-FKBP-Inp54 plus the plasma membrane-localizing Lyn_11_-FKB construct. Addition of rapamycin resulted in the rapid recruitment of Inp54 to the plasma membrane (Fig 6A, insets). As expected, macropinocytosis proceeded normally in cells expressing Rab5A, regardless of whether Inp54 was recruited or not to the membrane (Fig 6A, top panels). However, while cytosolic Inp54 was not able to overcome the effect of dominant negative Rab5A on macropinocytosis, the phosphatase enabled TMR-D uptake when recruited to the membrane at the time of EGF stimulation (Fig 6A, bottom panels). Recruitment of Inp54 to the plasma membrane restored macropinocytosis to levels similar to that of cells expressing WT Rab5A (Fig 6B). Taken together, these findings confirm that the breakdown of PtdIns(4,5)P_2_ to PtdIns(4)P in the macropinocytic cup is critical for the completion of macropinocytosis. Active Rab5 plays an essential role in this process by recruiting to the macropinocytic cup 5-phosphatase activity required for sealing, exerted primarily by OCRL and Inpp5b that are recruited through the adaptor APPL1.

**Figure 6.**
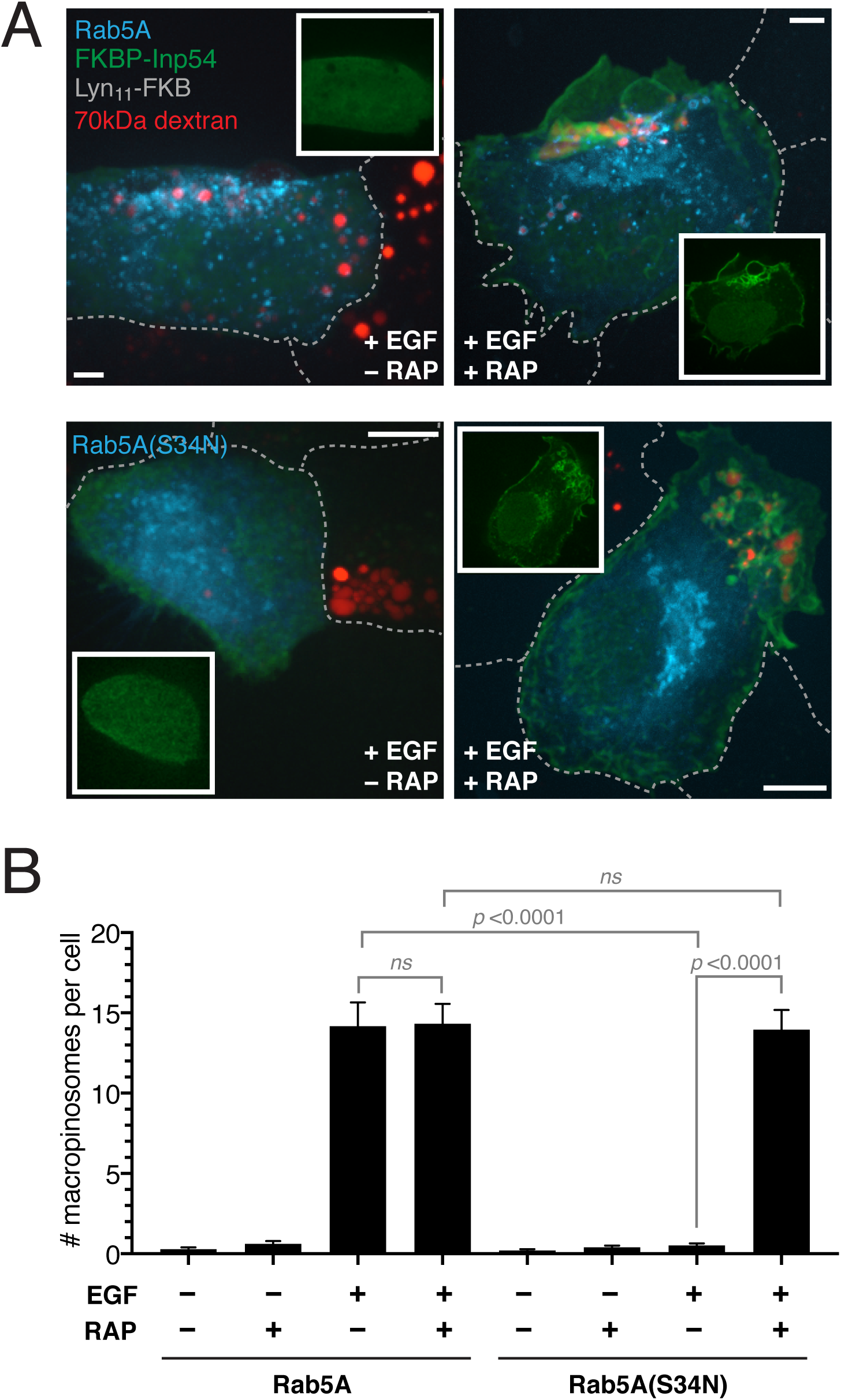
Depletion of PtdIns(4,5)P_2_ by recruitment of a 5-phosphatase overcomes the inhibitory effect of dominant negative Rab5A on macropinosome closure. **A)** Plasma membrane recruitment of Inp54 in Rab5A- or Rab5A_S34N_-expressing cells (cyan) was induced by rapamycin-dependent dimerization of FKBP-Inp54 (green) to Lyn_11_-FKB. Monolayers were treated with EGF ± rapamycin in the presence of TMR-D, as described in Materials and Methods. Following fixation, retention of TMR-D (red) was visualized in extended focus images. Insets: single *XY* slice illustrating the localization of Inp54. All scale bars: 5 μm. Images are representative of ≥ 30 fields from ≥ 3 separate experiments of each type. **B)** Effect of Inp54 recruitment on the number of macropinosomes formed per cell. Cells treated as in **A** were fixed, and the number of macropinosomes per cell counted by confocal microscopy. For each condition, 4 independent experiments were quantified, with ≥ 10 cells per replicate. Data are means ± SEM. *p* values were calculated using unpaired, 2-tailed student’s t-tests.

## Discussion

The involvement of Rab5 in macropinocytosis has been known for decades, but its mode of action remained enigmatic and controversial. Some reports concluded that Rab5 operated at the plasma membrane, promoting ruffling (Barbieri *et al*, 1998; Lanzetti *et al*, 2004; Spaargaren & Bos, 1999), while others disputed this claim and proposed instead a role for Rab5 only after vacuole internalization (Feliciano *et al*, 2011; Roberts *et al*, 2000). Here, we demonstrate that breakdown of PtdIns(4,5)P_2_ is a central requirement for macropinosome closure, a mechanism dependent on Rab5 and its effectors APPL1, Inpp5b and OCRL.

Our findings indicate that Rab5 exerts its effects prior to the scission of the nascent macropinocytic vacuole from the surface membrane. Accordingly, dominant-negative Rab5 prevents formation of the vacuole while not affecting the occurrence of membrane ruffling. The docking and likely fusion of Rab5-decorated vesicles with circular ruffles prior to sealing is consistent with this conclusion, as is the observation that the dominant-negative form of the GTPase is inhibitory when recruited to the plasmalemma. It is impossible using current technology to unambiguously ascertain that Rab5-positive vesicles in fact fuse with the surface membrane prior to scission, but experiments where fusion was precluded support this notion. Briefly, we implemented a new system whereby a dominant-negative form of NSF was acutely released into the cytosol of the test cells, obviating the detrimental non-specific effects associated with long-term expression of such inhibitory constructs. Blocking membrane fusion using this comparatively rapid maneuver caused a profound inhibition of macropinocytosis. Interestingly, the requirement for focal fusion of endomembranes with the plasma membrane during macropinocytosis is analogous to what has been reported for phagocytosis (Bajno *et al*, 2000; Braun *et al*, 2004).

The sealing of macropinosomes is accompanied by loss of PtdIns(4,5)P_2_ from their limiting membrane (Porat-Shliom *et al*, 2008; Welliver & Swanson, 2012), an observation we were able to verify (Fig 4). While firmly documented in the past, the functional implications of the disappearance of the inositide were unclear. Our data not only showed a clear correlation between PtdIns(4,5)P_2_ depletion and macropinosome sealing but, more pertinently, indicated that hydrolysis of the inositide on circular ruffles failed to occur when Rab5 activity was impaired (Fig 4B). Moreover, preventing the depletion of PtdIns(4,5)P_2_ by enhancing its rate of synthesis –through overexpression of PIP5K– precluded macropinosome formation. These findings led us to postulate that PtdIns(4,5)P_2_ hydrolysis from forming macropinosomes is essential for their scission from the membrane. Rapid PtdIns(4,5)P_2_ hydrolysis accompanies endosome and phagosome formation also, and may be a common feature and requirement of all endocytic pathways.

We noted that docking of Rab5 to circular ruffles preceded PtdIns(4,5)P_2_ depletion from the macropinocytic cup (Fig 4A) and therefore postulated a role for the GTPase in the hydrolysis of the phosphoinositide. Our subsequent experiments provided a plausible mechanism: we propose that through the adaptor APPL1, Rab5 recruits to the membrane Inpp5b and OCRL, two active phosphatases capable of degrading PtdIns(4,5)P_2_ to PtdIns(4)P. The adaptor, as well as the phosphatases, is recruited to nascent macropinosomes (Fig 5A-B) and, more importantly, their depletion greatly inhibits macropinocytosis, without altering membrane ruffling (Fig 5C-I).

Dephosphorylation by Inpp5b and OCRL is not solely responsible for the elimination of PtdIns(4,5)P_2_ from macropinosomes. Welliver and Swanson (2012) and Yoshida et al. (2015) had clearly demonstrated earlier that phospholipase C contributes to the process, generating diacylglycerol and Ins(1,4,5)P_3_. We believe that the initial hydrolysis of PtdIns(4,5)P_2_ is caused by phospholipase C, while the final stages of its removal, which occur at the time of sealing are catalyzed by Inpp5b and OCRL. Alternatively, phospholipase C activity may predominate at the base of macropinocytic cup, while the phosphatases may be most active at the singular point where sealing occurs. According to this scenario, restoration of inositol phosphatase activity should bypass the inhibition of macropinocytosis by dominant-negative Rab5. Indeed, using an inducible heterodimerization system we found that macropinocytosis could be restored in cells expressing Rab5A_S34N_ by recruiting an exogenous phosphatase, Inp54, to the membrane at the time of stimulation with EGF (Fig 6A-B).

In summary, our experiments have defined the mechanism by which Rab5 and its associated inositol 5-phosphatases OCRL and Inpp5b catalyze macropinosome sealing. The hydrolysis of PtdIns(4,5)P_2_ occurs in the macropinocytic cup as a result of the recruitment and eventual fusion of early endosomes containing Rab5 and its effectors. These data serve to highlight the intricate intersection between cytoskeletal dynamics, small GTPase activity and phosphoinositide signalling that occurs during dynamic cellular uptake processes like macropinocytosis.

## Materials and Methods

### Reagents

Mammalian expression vectors were obtained from the following sources: GFP-Rab5A (Roberts *et al*, 2000), GFP-Rab5A_S34N_ (plasmid #35141; Addgene), CFP-Rab5A (Heo *et al*, 2006), CFP-Rab5A_S34N_ (Heo *et al*, 2006), RFP-Rab5A (Chen *et al*, 2009), mCherry-Rab5A_S34N_ (plasmid #35139; Addgene), PLCδ-PH-GFP (Stauffer *et al*, 1998), PLCδ-PH-RFP (Stauffer *et al*, 1998), Lyn_11_-RFP (Lee *et al*, 2007), GFP-PIP5Kα (Fairn *et al*, 2009), GFP-PIP5Kβ (Fairn *et al*, 2009), GFP-PIP5Kβ_KD_ (Fairn *et al*, 2009), pC_4_S_1_-EGFP-F_M_4-FCS-hGH (Ariad Pharmaceuticals; (Rivera *et al*, 2000)), NSF_E329Q_ (Coppolino *et al*, 2002), GFP-APPL1 (Zoncu *et al*, 2009), Inpp5b-GFP (Williams *et al*, 2007), OCRL-GFP (Swan *et al*, 2010), Lyn_11_-FKB (plasmid #20147; Addgene), CFP-FKBP (plasmid #20160; Addgene) YF-FKBP-Inp54 (Suh *et al*, 2006).

Primary antibodies were purchased from the following vendors: Rab5 (catalogue #3547; Cell Signaling Technology), α-tubulin (catalogue #T5168; Sigma-Aldrich), GM130 (catalogue # 610822; BD Pharmingen). HRP, Alexa-568- and Alexa-647-conjugated secondary antibodies against mouse IgG were from Jackson ImmunoResearch Labs.

### Cell culture

The A431 cell line was obtained from and authenticated by the American Type Culture Collection (ATCC). Prior to experimentation, this cell line was revalidated by assessing expression of epithelial membrane markers (E-cadherin, β-catenin), and responsiveness to epidermal growth factor (EGF). A431 cells were grown in DMEM containing L-glutamine (MultiCell, Wisent) and 10% heat-inactivated FCS, at 37°C under 5% CO_2_. The cell line tested negative for mycoplasma contamination by DAPI staining.

### Macropinocytosis assays

A431 cells were plated on 18 mm glass coverslips at a concentration of 1.5 x 10^5^ cells mL^-1^ in DMEM containing L-glutamine and 10% heat-inactivated FCS, at 37°C under 5% CO_2_. The next day, medium was changed to HBSS medium (cat# 311-515-CL; MultiCell, Wisent), and cells were serum starved for 3 hr at 37°C. Cells were treated 20 min with 200 ng mL^-1^ EGF (Invitrogen, Thermo Fisher Scientific), in the presence of 100 μg mL^-1^ tetramethylrhodamine-conjugated 70 kDa dextran (TMR-D; Molecular Probes, Thermo Fisher Scientific). Monolayers were rinsed 3 times with PBS, placed in HBSS, and imaged live. The cells were visualized at the indicated intervals after addition of EGF ± TMR-D addition (5, 10, 15, 20 min).

To distinguish sealed macropinosomes from open circular ruffles, A431 cells were serum starved for 3 hr at 37°C and incubated 2 min with 200 ng mL^-1^ EGF and 100 μg mL^-1^ TMR-D. Cells were then washed 3 times with cold 1x PBS, and stained for 1 min with 20 μM FM4-64 (Invitrogen, Thermo Fisher Scientific) at 10°C. Samples were viewed immediately by confocal microscopy.

### DNA transfection

For transient transfection, A431 cells were plated on 18 mm glass coverslips at 1.5 x 10^5^ cells mL^-1^, 16-24 hr prior to experiments. Lipofectamine LTX with PLUS reagent (Invitrogen, Thermo Fisher Scientific) was used according to the manufacturer’s instructions. A431 cells were transfected at a 3:1 ratio using 1.5 μL Lipofectamine LTX, 0.5 μg DNA and 0.5 μL of PLUS reagent per well, for 4 hr in Opti-MEM medium (Gibco, Thermo Fisher Scientific). Following this, medium was changed to DMEM containing L-glutamine and 10% heat-inactivated FCS, and cells used for experiments 16 hr after transfection.

### Small interfering RNA gene silencing

siRNAs for Rab5 isoforms A, B, and C were from Sigma, based on previously described sequences (Huang *et al*, 2004). ON-TARGETplus human siRNA SMARTpools targeting APPL1 (cat# L-005138-00-0005), Inpp5b (cat# L-021811-00-0005) and OCRL (cat# L-01-0026-00-005) were obtained from Dharmacon, along with an ON-TARGETplus non-targeting pool (cat# D-001810-10-20). siRNA delivery was performed using either Nucleofector Kit V (Lonza) or Neon transfection system (Life Technologies, Thermo Fisher Scientific), according to the manufacturers’ protocols. For nucleofection of Rab5a/b/c siRNA, 1 x 10^6^ cells A431 cells were resuspended in 100 μL of Nucleofector Solution T, containing 200 pmol of Rab5a/b/c siRNA, and electroporated with program X-01 on the Nucleofector I system (Lonza). Cells were used 48 h after siRNA treatment. For Neon transfection of APPL1/Inpp5b/OCRL siRNA smart-pools, A431 cells were resuspended to 5 x 10^6^ cells mL^-1^, and 100 μL of suspension mixed with 200 pmol siRNA. Electroporation was then done using two 20 ms pulses of 1400 V. After electroporation, cells were immediately transferred to DMEM containing L-glutamine and 10% heat-inactivated FCS, before seeding on coverslips at concentration of 2.5 x 10^6^ cells mL^-1^. siRNA-treated cells were used for macropinocytosis experiments 48 hr after electroporation. In some cases, after 24 hr, cells were transiently transfected with mammalian expression vectors as described above.

siRNA-mediated knockdown was confirmed at 48 hr by either Western blotting or quantitative PCR. For Rab5A/B/C knockdown, cells were lysed in Laemmli buffer (Bio-Rad). Lysates were separated by SDS-PAGE, followed by transfer to a polyvinylidene difluoride membrane. The membrane was blocked in TBS containing 5% BSA and 0.05% Tween-20 for 30 min at room temperature, followed by primary antibody staining for 1 hr at room temperature, in blocking buffer. Primary antibody dilutions: Rab5 (1:1000), α-tubulin (1:1000; loading control). After washing the membrane in TBS containing 0.05% Tween-20, samples were incubated 30 min at room temperature with an HRP-conjugated secondary antibody at 1:5000 dilution. Blots were visualized using the ECL Prime Western Blot detection reagent (GE Healthcare) on film.

For validation of APPL1, Inpp5b and OCRL knockdown, RNA was isolated from siRNA-treated cells using the GeneJET RNA Purification kit (Thermo Fisher Scientific). 1 µg of RNA was used for single strand cDNA synthesis with the SuperScript VILO cDNA Synthesis kit (Invitrogen, Thermo Fisher Scientific). Quantitative PCR was done using the TaqMan System (Applied Biosystems, Thermo Fisher Scientific) and TaqMan Fast Advanced Mastermix (Applied Biosystems, Thermo Fisher Scientific) on a Step One Plus Real-Time PCR thermal cycler (Step One software v2.2.2; Applied Biosystems, Thermo Fisher Scientific). The Taqman gene expression assays for the reference gene (GAPDH – ID# Hs02786624_g1) and target genes (APPL1 – ID# Hs00179382_m1; Inpp5b – ID# Hs00299982_m1; OCRL – ID# Hs00240844_m1) were duplexed in triplicate. Target gene expression was determined by quantification relative to the GAPDH reference gene and the non-targeting pool control sample (ΔΔ*C*_*T*_ method; (Livak & Schmittgen, 2001)).

### Rapamycin-inducible system for targeted protein translocation

For Rab5 iRAP, full length human Rab5A or Rab5A(S34N) were excised from GFP expression vectors using HindIII/ BamHI or XhoI/BamHI sites, respectively. These fragments were cloned downstream of CFP-FKBP using the same sites, and co-expressed in cells with the Lyn_11_-FRB plasma membrane localizing sequence. Treatment of A431 cells with 2.5 μM rapamycin mediated the rapid translocation of FKBP-CFP-Rab5A/Rab5A(S34N) to the plasma membrane by induced heterologous dimerization of rapamycin-binding FKBP and FKB domains. After 5 min pre-treatment with rapamycin, macropinocytosis assays (TMR-D uptake) were performed as described above, in the presence of rapamycin. iRAP experiments using YFP-FKBP-Inp54 and Lyn_11_-FRB, to recruit Inp54 to the plasma membrane upon addition of rapamycin were done similarly.

### *In vitro* reverse dimerization system

pC_4_S_1_-EGFP-F_M_4-FCS-hGH is a part of an *in vitro* reverse dimerization system (Ariad Pharmaceuticals; (Rivera *et al*, 2000)). This vector contains a CMV enhancer/promoter (C), the secretion signal sequence of human growth hormone (S), EGFP, 4 tandem repeats of rapalog-binding FKBP_36M_ (F_M_4), and the human growth hormone coding sequence (hGH) downstream of a furin cleavage signal (FCS). To create a reverse dimerization construct for NSF_E329Q_, the pC_4_S_1_-EGFP-F_M_4-FCS-hGH vector was modified by cloning. The hGH signal sequence was removed using EcoRI/XbaI digestion, and replaced with a synthetic Kozak sequence. Next, the FCS-hGH SpeI/BamHI fragment was replaced with NSF_E329Q_. The resultant vector, pC_4_-EGFP-F_M_4-NSF_E329Q_, generates a protein that undergoes spontaneous aggregation that reverses upon the addition of D/D solubilizer (catalogue# 635054; Takara Bio), a cell-permeant rapamycin analog that binds to the F_M_4 domains (see Fig 3e).

A431 cells were transiently transfected with pC_4_-EGFP-F_M_4-NSF_E329Q_, as described above. Transfected monolayers were then serum-starved in HBSS for 2 h, the last hour of which included treatment with 500 nM D/D solubilizer or vehicle (DMSO) control. Macropinocytosis assays (i.e. uptake of TMR-D) were performed as above. The reverse dimerization of NSF_E329Q_ was confirmed in transfected monolayers by assessment of the effect of pC_4_-EGFP-F_M_4-NSF_E329Q_ expression ± D/D solubilizer on Golgi morphology, using immunostaining. Briefly, 1 hr after treatment with 500 nM D/D solubilizer or vehicle (DMSO) control, transfected monolayers were fixed in 3% paraformaldehyde for 10 min at room temperature. Monolayers were blocked/permeabilized in PBS containing 5% BSA and 0.1% saponin, and stained with antibody to GM130 (1:500 dilution in blocking buffer), followed by fluorescently-conjugated secondary antibody.

### Confocal microscopy and image analysis

Confocal images were acquired using a Yokogawa CSU10 spinning disk system (Quorum Technologies Inc.). Images were acquired using a 63×/1.4 NA oil objective or a 25x/0.8 NA water objective (Zeiss), as indicated, with an additional 1.5x magnifying lens. For live experiments, cells were maintained at 37°C using an environmental chamber (Live Cell Instruments). Routine analyses were done using Volocity software (Perkin Elmer) or Fiji (Schindelin *et al*, 2012).

For colocalization analyses, Volocity software was used to calculate positive product of the differences of the mean (PDM) channels (Li et al., 2004), which were then overlaid on merged images for visualization. For fluorescence intensity calculations, background-subtracted intensities per unit area for expressed fluorescent protein constructs or endogenous proteins (immunofluorescence) were measured in Volocity software. Ratios were calculated comparing relative intensities of transfected markers in the macropinosome membrane compared to plasma membrane, as indicated in the text. All statistics were calculated using GraphPad Prism software (GraphPad Software, Inc.).

For the quantitation of ruffling, a Java plug-in (Ruffle_Analysis.java) was written for Fiji to analyze background-subtracted ruffling movies of EGF-treated A431 cells expressing Lyn_11_-RFP. Movies were recorded for 2 min pre-EGF and 8 min post-EGF treatment (200 ng mL^-1^). For a selected region of the ruffling plasma membrane, the plug-in calculates a summed squared pixel difference within a user defined ROI, between frame_*n*+*1*_ and frame_*n*_, for every frame pair in a background subtracted time-lapse (background subtracted ΣΔ_pixel_^2^). The code first takes each individual pixel value within a square/rectangular ROI and subtracts it from the corresponding pixel value in the next frame (Δ_pixel_). Next, the differences are squared (Δ_pixel_^2^) to eliminate the cancelling effects of positive vs. negative changes in fluorescence intensity. Finally, these values are then summed generating a (ΣΔ_pixel_^2^) value for each frame, starting with frame_*n*+*1*_. For normalization, background-subtracted ΣΔ_pixel_^2^ values were normalized to the mean pre-EGF values. For each cell, the standard deviation of the background-subtracted ΣΔ_pixel_^2^ for the pre-EGF period was compared to that of the post-EGF period, as a metric of the continuous intensity changes/pixel that are seen in an ROI during plasma membrane ruffling. Additionally, temporal projections of A431 cells pre- and post-EGF treatment were generated in Fiji, using the included Temporal Color Code plug-in (Daste *et al*, 2017) and cool LUT. Data calculations and normalizations were done using Microsoft Excel software (Microsoft Corporation). All statistics were calculated using GraphPad Prism software (GraphPad Software, Inc.).

## Author contributions

MM, HS, AV, JB, SG designed research; MM, HS, AV performed experiments; MM quantified data; MM, SG prepared figures; MM, SG wrote manuscript; MM, HS, AV, JB, SG reviewed draft; SG provided funding. All authors have read and approve of the final manuscript.

## Acknowledgments

We thank Dr. Philip Ostrowski for writing the ruffle quantitation Fiji plugin, and Dr. Michal Bohdanowicz for construction of Rab5A and Rab5A_S34N_ iRAP constructs. The plasmid for the reverse dimerization system was kindly provided by Dr. Victor Rivera (Ariad Pharmaceuticals). MM was the recipient of a Heart and Stroke Pfizer Research Fellowship. Work in the authors’ laboratory is supported by Canadian Institutes of Health Research grant FDN-143202.

## Data availability

The data that support the findings of this study are available from the corresponding author upon reasonable request.

## Code availability

Ruffle_Analysis.java plugin for ImageJ/FIJI is available at: https://doi.org/10.6084/m9.Fighare.12349967

The authors declare no conflict of interest.

## Appendix Figure Legends

**Appendix Figure S1.**
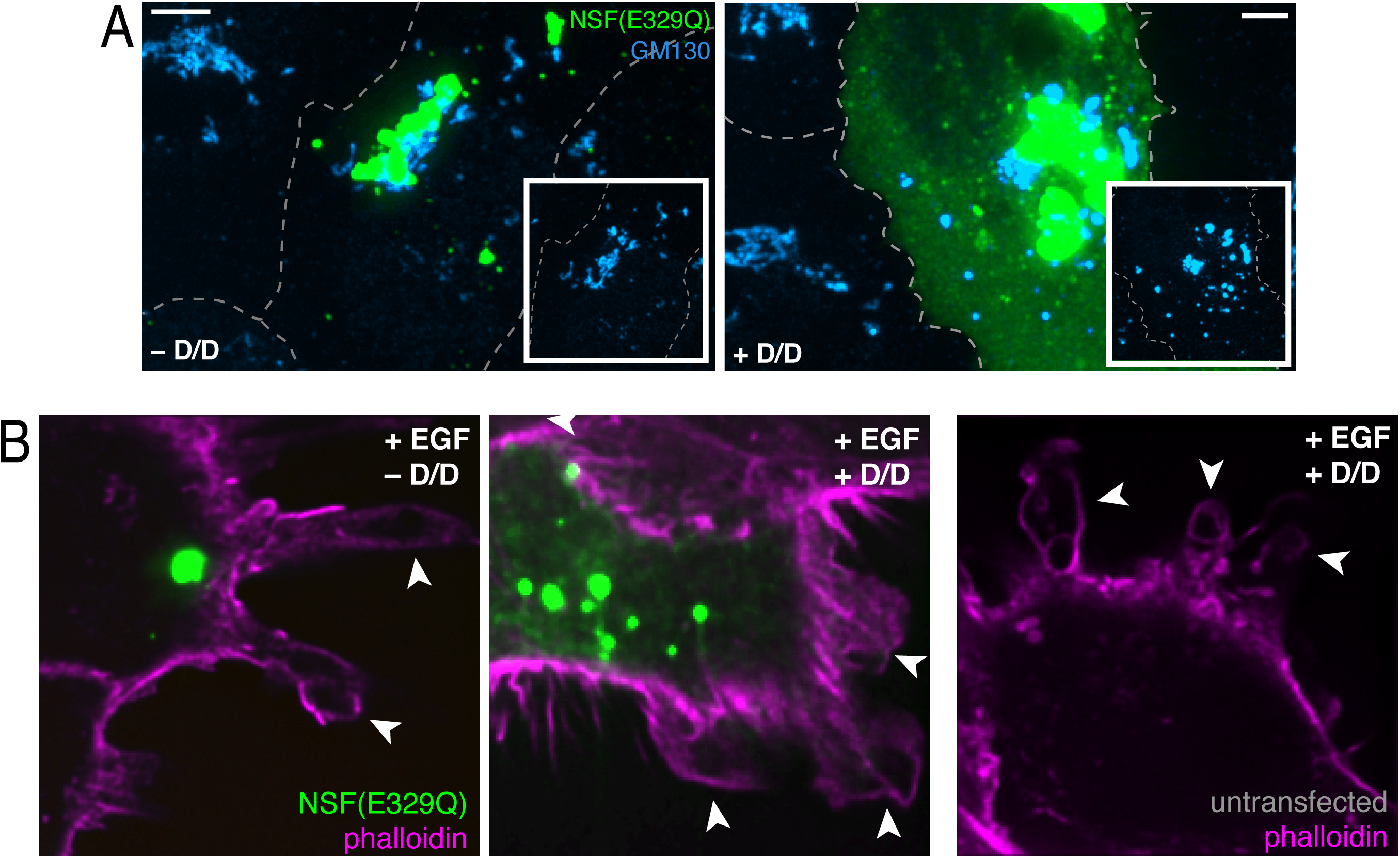
**A)** Cells expressing EGFP-F_M_4-NSF_E329Q_ were incubated for 60 min without (left) of with D/D solubilizer (right). Following fixation, cells were monitored for aggregation of NSF_E329Q_ (green) and immunostained for GM130 (cyan). Insets: GM130 immunostain alone. **B)** Cells expressing EGFP-F_M_4-NSF_E329Q_ were treated for 60 min ± D/D solubilizer, as indicated, followed by EGF, then fixed and permeabilized. Cells were stained for phalloidin to visualize actin-rich ruffles (magenta). Untransfected control cells that had been treated with D/D solubilizer and EGF shown for reference at right. All scale bars: 5 μm. Images are representative of ≥ 30 fields from ≥ 3 separate experiments of each type.

**Appendix Figure S2.**
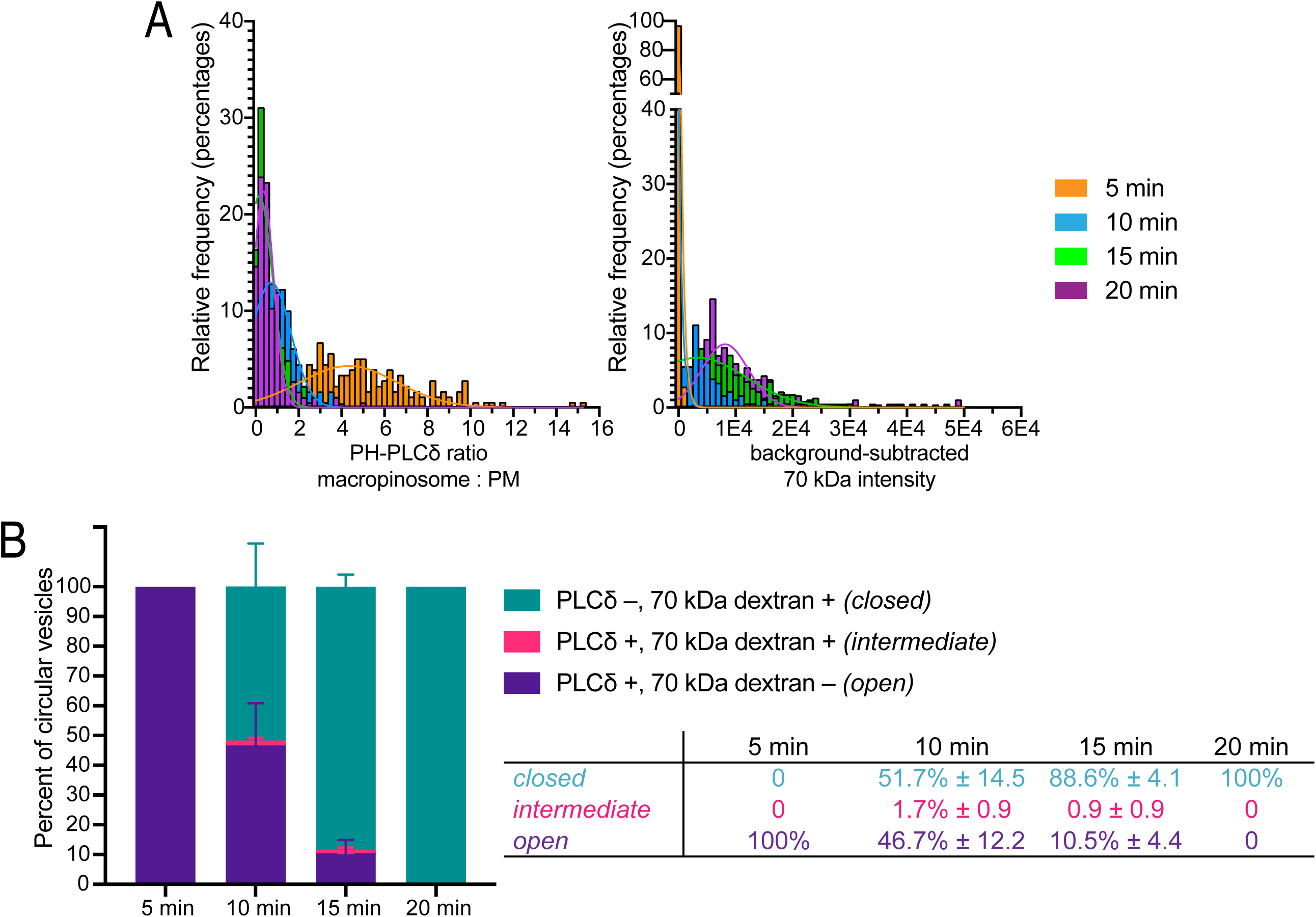
**A)** Frequency histograms of PLCδ-PH-GFP macropinosome:plasma membrane ratio (left) and background-subtracted TMR-D intensity (right) at: 5 min (orange), 10 min (blue), 15 min (green), and 20 min (magenta) after stimulation with EGF. **B)** Analysis of the percentage of closed, intermediate or open vacuoles, at 5, 10, 15 and 20 min after EGF addition. Degree of closure was judged based on their PLCδ-PH-GFP content (normalized per unit plasma membrane) and TMR-D content. For each condition, 3 independent experiments were quantified, with ≥ 50 vesicles per replicate. The numerical percentages are included at right. Data are means ± SEM.

**Appendix Figure S3.**
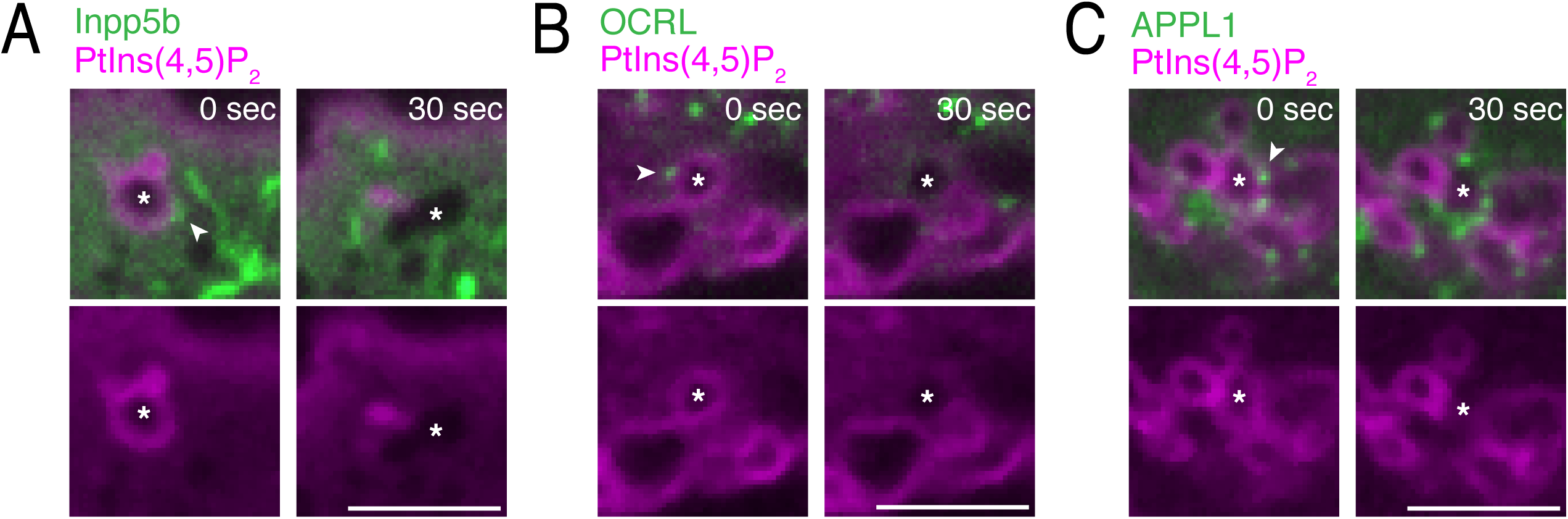
Live cell visualization of PtdIns(4,5)P_2_ during EGF-induced macropinocytosis using PLCδ-PH-RFP (magenta) in cells co-expressing GFP-tagged-**A)** Inpp5b, **B)** OCRL, or **C)** APPL1. Asterisks mark macropinocytic cups of interest. Arrowheads indicate the apposition of GFP-positive vesicles to PtdIns(4,5)P_2_-positive macropinocytic cups (left panels), that preceded PtdIns(4,5)P_2_ depletion detected 30 sec later (right panels). All scale bars: 5 μm. Images are representative of ≥ 20 fields from ≥ 2 separate experiments of each type.

**Appendix Movie 1.** Corresponding to Fig 1D. Cells expressing Lyn_11_-RFP (white) and GFP-Rab5A (not shown here, see Fig 1D) imaged for plasma membrane ruffling every 20 sec during EGF treatment. EGF addition occurred between 120-140 sec timestamps, as indicated. Acquisition rate: 6 fps. Scale bar: 5 μm. Movie is representative of ≥20 movies from 4 separate experiments.

**Appendix Movie 2.** Corresponding to Fig 1F. Cells expressing Lyn_11_-RFP (white) and GFP-Rab5A_S34N_ (not shown here, see Fig 1F) imaged for plasma membrane ruffling every 20 sec during EGF treatment. EGF addition occurred between 100-120 sec timestamps, as indicated. Acquisition rate: 6 fps. Scale bar: 5 μm. Movie is representative of ≥20 movies from 4 separate experiments.

**Appendix Movie 3.** Corresponding to Fig 2A and B. Cells expressing Lyn_11_-RFP (magenta) and GFP-Rab5A (cyan) imaged every 20 sec during EGF treatment. EGF addition occurred between 100-120 sec timestamps, as indicated. Arrowhead follows a macropinosome of interest. Acquisition rate: 4 fps. Scale bar: 5 μm. Movie is representative of ≥20 movies from 3 separate experiments.

**Appendix Movie 4.** Corresponding to Fig 4C. Cells expressing Lyn_11_-RFP (magenta) and PLCδ-PH-GFP (green) imaged every 30 sec during EGF treatment. Colocalization of PLCδ-PH-GFP with Lyn_11_-RFP shown in yellow. EGF addition occurred between 150-180 sec timestamps, as indicated. Arrowhead follows a macropinosome of interest. Acquisition rate: 4 fps. Scale bar: 5 μm. Movie is representative of ≥20 movies from 3 separate experiments.

**Appendix Movie 5.** Corresponding to Fig 5E. Scrambled siRNA-treated cells expressing Lyn_11_-RFP (white) imaged for plasma membrane ruffling every 20 sec during EGF treatment. EGF addition occurred between 120-140 sec timestamps, as indicated. Acquisition rate: 6 fps. Scale bar: 5 μm. Movie is representative of ≥30 movies from 4 separate experiments.

**Appendix Movie 6.** Corresponding to Fig 5G. APPL1 siRNA-treated cells expressing Lyn_11_-RFP (white) imaged for plasma membrane ruffling every 20 sec during EGF treatment. EGF addition occurred between 120-140 sec timestamps, as indicated. Acquisition rate: 6 fps. Scale bar: 5 μm. Movie is representative of ≥30 movies from 4 separate experiments.

